# Pathogenic amino acids in mitochondrial proteins more frequently arise in lineages closely related to human than in distant lineages

**DOI:** 10.1101/119792

**Authors:** Galya V. Klink, Andrey V. Golovin, Georgii A. Bazykin

## Abstract

Propensities for different amino acids within a protein site change in the course of evolution, so that an amino acid deleterious in a particular species may be acceptable at the same site in a different species. Here, we study the amino acid-changing variants in human mitochondrial genes, and analyze their occurrence in non-human species. We show that substitutions giving rise to the human amino acid variant tend to occur in lineages closely related to human more frequently than in more distantly related lineages, indicating that a human variant is more likely to be deleterious in more distant species. Unexpectedly, amino acids corresponding to pathogenic alleles in humans also more frequently originate at more closely related lineages. Therefore, a pathogenic variant still tends to be more acceptable in human mitochondria than a variant that may only be fit after a substantial perturbation of the protein structure.

**Significance:** Homologous proteins can carry different amino acids at the same position in different species. These changes can be neutral, or can reflect differences in the pressure of selection on these sites. We hypothesized that amino acids observable in human population appear on average more frequently in close than in distant human relatives. For mitochondrial proteins, we observe this for both frequent and rare human alleles. Unexpectedly, substitutions that would be pathogenic in humans also more frequently appear in species more closely related to humans than in distantly related ones. Therefore, despite their pathogenicity, these variants are on average more acceptable in humans than other amino acids that were observed at this site in distantly related species.

## Introduction

Fitness conferred by a particular allele depends on a multitude of factors, both internal to the organism and external to it. Therefore, relative preferences for different alleles change in the course of evolution due to changes in interacting loci or in the environment. In particular, changes in the propensities for different amino acid residues at a particular protein position, or single-position fitness landscape, have been detected using multiple approaches (SPFL, Bazykin, 2015; Harpak, Bhaskar, & Pritchard, 2016; Storz, 2016).

One way to observe such changes is by analyzing how amino acid variants (alleles) and substitutions giving rise to them are distributed over the phylogenetic tree. In particular, multiple substitutions giving rise to the same allele, or homoplasies, are more frequent in closely related than in distantly related species – a pattern expected if frequent homoplasies mark the segments of the phylogenetic tree where the arising allele confers high fitness (Goldstein, Pollard, Shah, & Pollock, 2015; Naumenko, Kondrashov, & Bazykin, 2012; Povolotskaya & Kondrashov, 2010; Rogozin, Thomson, Csürös, Carmel, & Koonin, 2008; Zou & Zhang, 2015).

Another manifestation of changes in SPFL is the fact that amino acids deleterious and, in particular, pathogenic in humans are often fixed as the wild type in other species. This phenomenon has been termed compensated pathogenic deviations, under the assumption that human pathogenic variants are non-pathogenic in other species due to compensatory (or permissive) changes elsewhere in the genome (Jordan et al., 2015; Kondrashov, Sunyaev, & Kondrashov, 2002; Soylemez & Kondrashov, 2012). The distribution of the evolutionary distances to the nearest vertebrate species in which a human pathogenic variant is observed in the nuclear genome is well approximated by a sum of two exponential distributions, suggesting that just one compensatory change is typically required to permit a formerly deleterious variant (Jordan et al., 2015).

SPFLs of mitochondrial protein-coding genes also change with time (Goldstein et al., 2015; Klink & Bazykin, 2017; Zou & Zhang, 2015), and some of these changes may be due to intragenic or intergenic epistasis (Ji et al., 2014; Xie et al., 2016). Here, we use our previously developed approach for the study of phylogenetic clustering of homoplasies at individual protein sites (Klink & Bazykin, 2017) to ask how the fitness conferred by the amino acid variants observed in human mitochondrial proteins, either as benign or damaging, changes with phylogenetic distance from the human.

## Materials and Methods

### Data

Here, we reanalyzed the phylogenetic data on a set of 5 mitochondrial protein-coding genes in 4350 species of opisthokonts (Klink & Bazykin, 2017), as well as on each of the 12 mitochondrial protein-coding genes in several thousand species of metazoans, focusing on the amino acids present in the human mitochondrial genome.

A joint alignment of five concatenated mitochondrial genes of opisthokonts and alignments of 12 mitochondrial proteins of metazoans (Breen et al. 2012) were obtained as described in (Klink & Bazykin, 2017). These alignments were used to reconstruct constrained phylogenetic trees, ancestral states and phylogenetic positions of substitutions (Klink & Bazykin, 2017). As the reference human allele, we used the revised Cambridge Reference Sequence (rCRS) of the human mitochondrial DNA (Andrews et al., 1999). As non-reference alleles, we used amino acid changing variants from “mtDNA Coding Region & RNA Sequence Variants” section of the MITOMAP database. As pathogenic alleles, we used amino acid changing variants with “reported” (i.e., supported by one publication) or “confirmed” (i.e., supported by at least two independent publications) status from the “Reported Mitochondrial DNA Base Substitution Diseases: Coding and Control Region Point Mutations” section of MITOMAP.

### Clustering of substitutions giving rise to the human amino acid around the human branch

For each amino acid site in a protein, we considered those amino acid variants that (i) constitute the reference allele in humans and had arisen in the human lineage at some point during its evolution, or (ii) had originated in humans as a derived polymorphic allele, or (iii) are annotated in humans as pathogenic alleles. For further consideration, we retained only such alleles from each class for which at least one homoplasic (i.e., giving rise to the same allele by way of parallelism, convergence, or reversal) and at least one divergent (i.e., giving rise to a different allele) substitution from the same ancestral variant was observed at this site elsewhere on the phylogeny outside of the human lineage. Substitutions, including reversals, that occurred anywhere on the path between the root and *H. sapiens* were excluded. While the homoplasic and divergent substitutions had to derive from the same ancestral variant, it could be either the same or a different variant than that ancestral to the variant observed in human.

For each such allele, we compared the phylogenetic distances between human and positions of homoplasic substitutions with the distances between human and positions of divergent substitutions, using a previously described procedure which controls for the differences in SPFLs between sites or in mutational probabilities of different substitutions (Klink & Bazykin, 2017). Briefly, for each ancestral amino acid at each site, we subsampled equal numbers of homoplasic substitutions to human (reference, non-reference or pathogenic) amino acids and divergent substitutions to non-human amino acids. We then pooled these values across all considered sites and groups of derived alleles. Next, we categorized them by the phylogenetic distance between the human and the position of the substitution, and calculated, for each bin of the phylogenetic distances, the ratio of the numbers of homoplasic (H) and divergent (D) substitutions (H/D). To obtain the mean values and 95% confidence intervals for the H/D statistic, we bootstrapped sites in 1000 replicates, each time repeating the entire resampling procedure. As a control, we performed the same analyses using instead of the human variant a random non-human amino acid among those observed at this site, or using data obtained by simulating the evolution at each site along the same phylogeny and with gene-specific GTR+Gamma amino acid substitution matrices (Klink & Bazykin, 2017).

### Molecular dynamics simulation of single point mutations in position 91 of COX3

The amino acid at position 91 of COX3 is adjacent to the pore in the complex IV of the respiratory chain which is thought to be required for the transport of oxygen. Therefore, estimation of the effect of the mutation in this position requires a complete description of the cytochrome c oxidase complex in membrane environment. We applied coarse grain description with martini forcefield (Marrink, Risselada, Yefimov, Tieleman, & de Vries, 2007). Human cytochrome C oxydase was modelled with homology modelling approach using Modeller 9.18 (Eswar et al., 2006). The membrane environment was rebuild from a random position of lipids and restrained protein structure as in MemProt MD database (Stansfeld et al., 2015). Each mutant was subjected to 0.5 mks simulation with two replicas in GROMACS 2016 (Abraham et al., 2015) with time step 10fs with PME electrostatics (Wennberg et al., 2015). Simulation results were analyzed with MDAnalysis Python module (Michaud-Agrawal, Denning, Woolf, & Beckstein, 2011). Water molecules within the pore were counted in a cylindrical selection with radius 2nm, height 5nm and center position determined dynamically as the center of mass of the two helices of the dimer harboring mutations. Counts of water molecules and their standard deviations were estimated from the last 100 ns of the trajectory.

## Results

### Substitutions giving rise to reference human amino acids are more frequent in species closely related to human

The phylogenetic distribution of homoplasic (parallel, convergent or reversing) substitutions is relevant for understanding which amino acids are permitted at a particular species, and which are selected against. In particular, an excess of such substitutions giving rise to a particular allele in closely related lineages implies that the fitness conferred by this allele in these species is higher than in other species. Here, using phylogenies reconstructed from two datasets of mitochondrial protein coding genes obtained earlier (Klink & Bazykin, 2017), we asked how homoplasic substitutions giving rise to the human variant are positioned phylogenetically relative to the human branch, compared with other (divergent) substitutions.

The reference human allele is also observed, on average, in over half of other considered species (71.2% of species of opisthokonts for the 5-genes dataset, and 60% of all species of metazoans for the 12-genes dataset). In 7.1% (10%) of these species, it did not share common ancestry with the human, but instead originated independently in an average of 30 (21) independent homoplasic substitutions per site (Supporting Information, Table S1 and Figures S1-S2).

We asked whether the phylogenetic positions of homoplasic substitutions giving rise to the human allele are biased, compared to the positions of divergent substitutions giving rise to other alleles. This analysis controls for the biases associated with pooling sites and amino acid variants (see Materials and Methods).

In most proteins, the mean phylogenetic distances from human to the branches at which the human reference amino acid emerged independently due to a homoplasy were ~10% shorter than to the branches at which another amino acid emerged (Fig 1a; SI, Figure S3). No such decrease was observed for a random amino acid among those that were observed at this site in non-human species, or in simulated data (figure 1a; SI, figure S3). The number of homplasic substitutions giving rise to the human reference amino acid relative to the number of divergent substitutions towards non-human amino acids (H/D ratio) uniformly decreases with the evolutionary distance from the human (Fig 1b; S4 Fig). At most genes, the relative number of homoplasic substitutions giving rise to the human allele drops 1.5-3-fold with phylogenetic distance from the human branch (Fig 1b; SI, figure S4).

**Figure 1.**
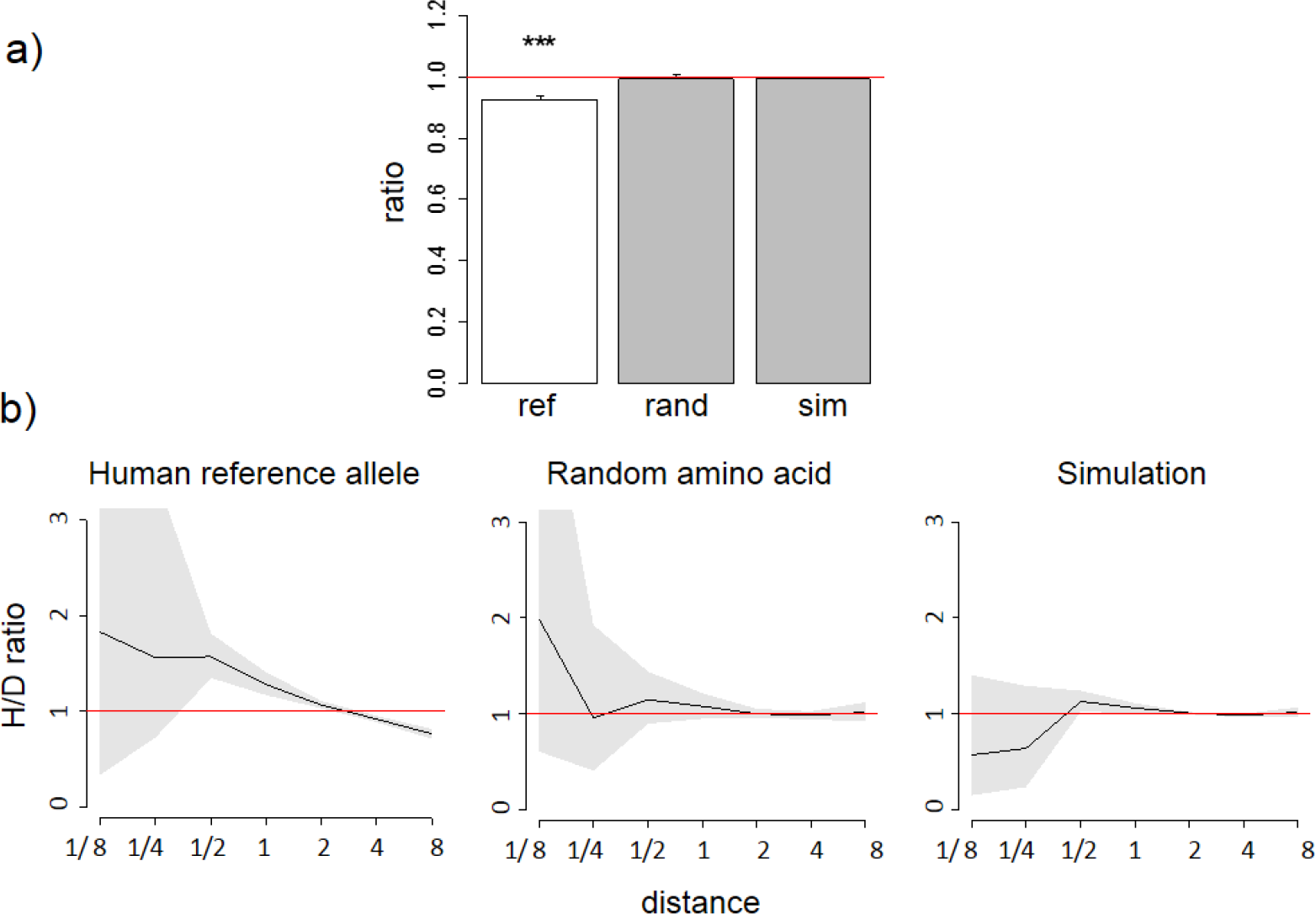
A reduced ratio of the phylogenetic distances between the human branch and substitutions to the considered amino acid vs. to other amino acids at the same site (a) and higher fraction of homoplasic substitutions to the human reference amino acid, compared with random amino acids that had independently originated at this site (H/D ratio) in species closely related to human (b) are observed for the human reference amino acid, but not for a random allele observed at this site or a simulated allele, in the 4350-species opisthokonts phylogeny. a) Ratios <1 imply that the considered allele arises independently closer at the phylogeny to humans than other alleles. The bar height and the error bars represent respectively the median and the 95% confidence intervals obtained from 1,000 bootstrap replicates, and asterisks show the significance of difference from the one-to-one ratio (*, P<0.05; **, P<0.01;***, P<0.001). ref, human reference allele; rand, a random non-human amino acid among those that were present in the site; sim, human allele in simulated data. b) Horizontal axis, distance between branches carrying the substitutions and the human branch, measured in numbers of amino acid substitutions per site, split into bins by log_2_(distance). Vertical axis, H/D ratios for substitutions at this distance. Black line, mean; grey confidence band, 95% confidence interval obtained from 1000 bootstrapping replicates. The red line shows the expected H/D ratio of 1. Arrows represent the distance between human and *Drosophila*.

The excess of homoplasic changes to the human reference amino acid at small phylogenetic distances from human is not an artefact of differences in mutation rates between amino acids in distinct clades, since these rates are similar in all considered species and cannot lead to such clustering (Klink & Bazykin, 2017). It is also not an artefact of differences in codon usage bias between species, as it was still observed when we considered only “accessible” amino acid pairs where the derived amino acid could be reached through a single nucleotide substitution from any codon of the ancestral amino acid (S3 Fig).

### Amino acids corresponding to variant alleles at human polymorphic sites more frequently arise in species closely related to humans

Next, we considered human SNPs in mitochondrial proteins in the MITOMAP database (Lott et al., 2013). We analyzed the phylogenetic distribution of substitutions in non-human species giving rise to the amino acid that is also observed as the non-reference (usually minor) allele in humans (SI, table S2).

Similarly to the human reference amino acid variant, the homoplasic substitutions giving rise to non-reference alleles were clustered on the phylogeny near humans, compared to the divergent substitutions giving rise to a variant never observed in humans (Fig 2; SI, figures S5-S6). Again, the mean phylogenetic distance from human to a substitution giving rise to the human non-reference amino acid was ~10% lower than to other substitutions (figure 2a; SI, figure S5), and the density of homoplasic substitutions giving rise to such an allele dropped significantly with phylogenetic distance from the human branch (Fig 2b; SI, figure S6). As before, this clustering was also observed if only accessible pairs of amino acids were considered, while no systematic differences were observed for random amino acids (SI, figure S5).

**Figure 2.**
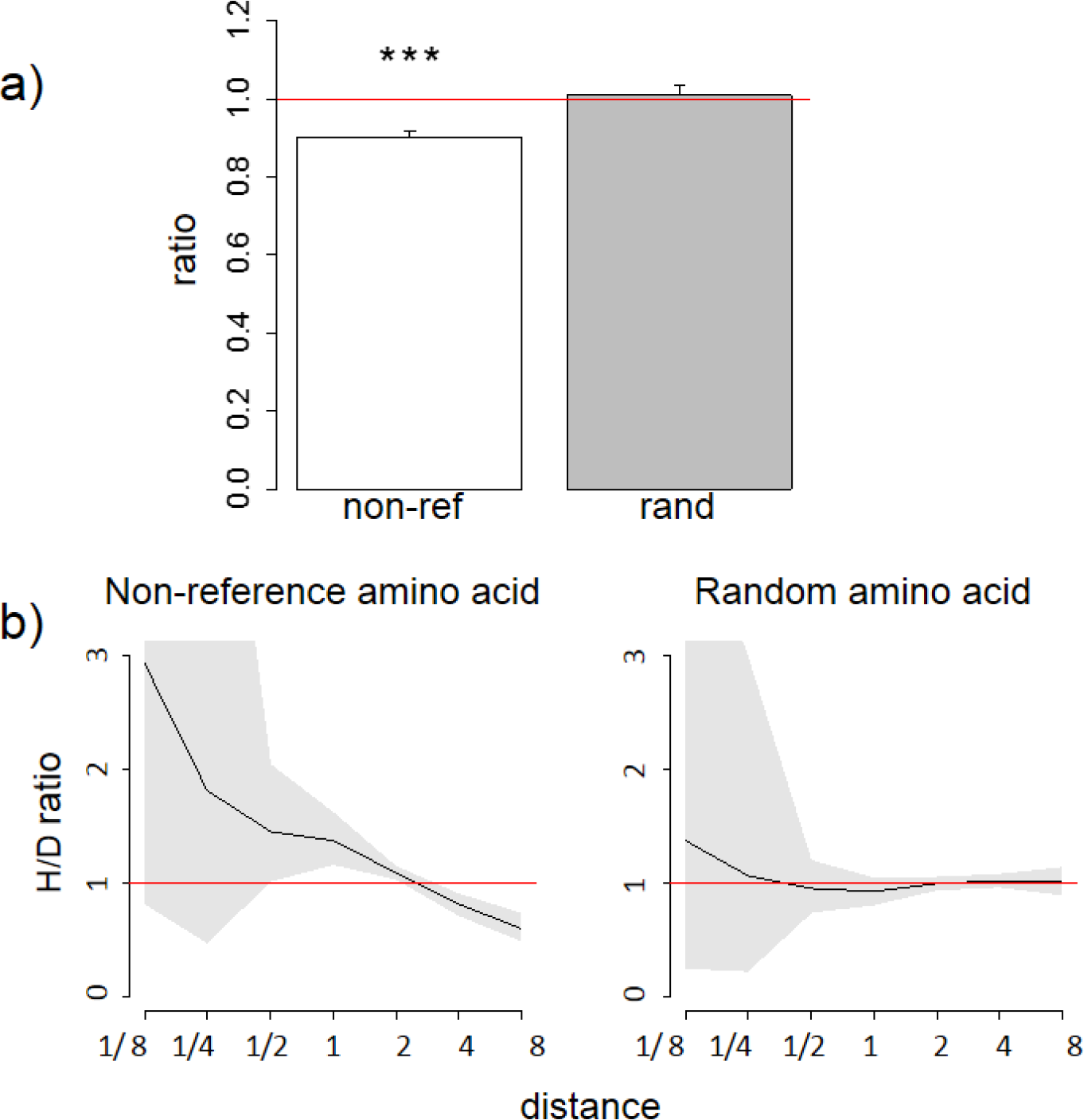
A reduced ratio of the phylogenetic distances between the human branch and substitutions to the considered amino acid vs. to other amino acids at the same site (a) and a higher H/D ratio in species closely related to human (b) are observed for the human non-reference amino acid, but not for a random allele observed at this site or a simulated allele, in the 4350-species opisthokonts phylogeny. Notations same as in Figures 1 and 2.

### Amino acids corresponding to human pathogenic variants more frequently arise in species closely related to humans

Finally, we considered human alleles annotated as disease-causing in the MITOMAP database (Lott et al., 2013). Since only a handful of mutations is thus annotated in each gene (SI, table S3), the variance in the estimates of the H/D ratio is, as expected, large. Still, in the opisthokont dataset (figure 3), as well as in five of the twelve genes of the metazoan dataset (SI, figures S7-S8), the human pathogenic variant also arose independently more frequently in the phylogenetic vicinity of humans. The opposite pattern, i.e., biased occurrence of the human pathogenic variant in phylogenetically remote species, has not been observed in any of the genes. This trend was even stronger for the six pathogenic mutations confirmed by two or more independent studies (“confirmed” status in MITOMAP; figure 3b). As before, this result is not due to preferential usage of codons more likely to mutate into the human variant in species closely related to human (SI, figure S7).

**Figure 3.**
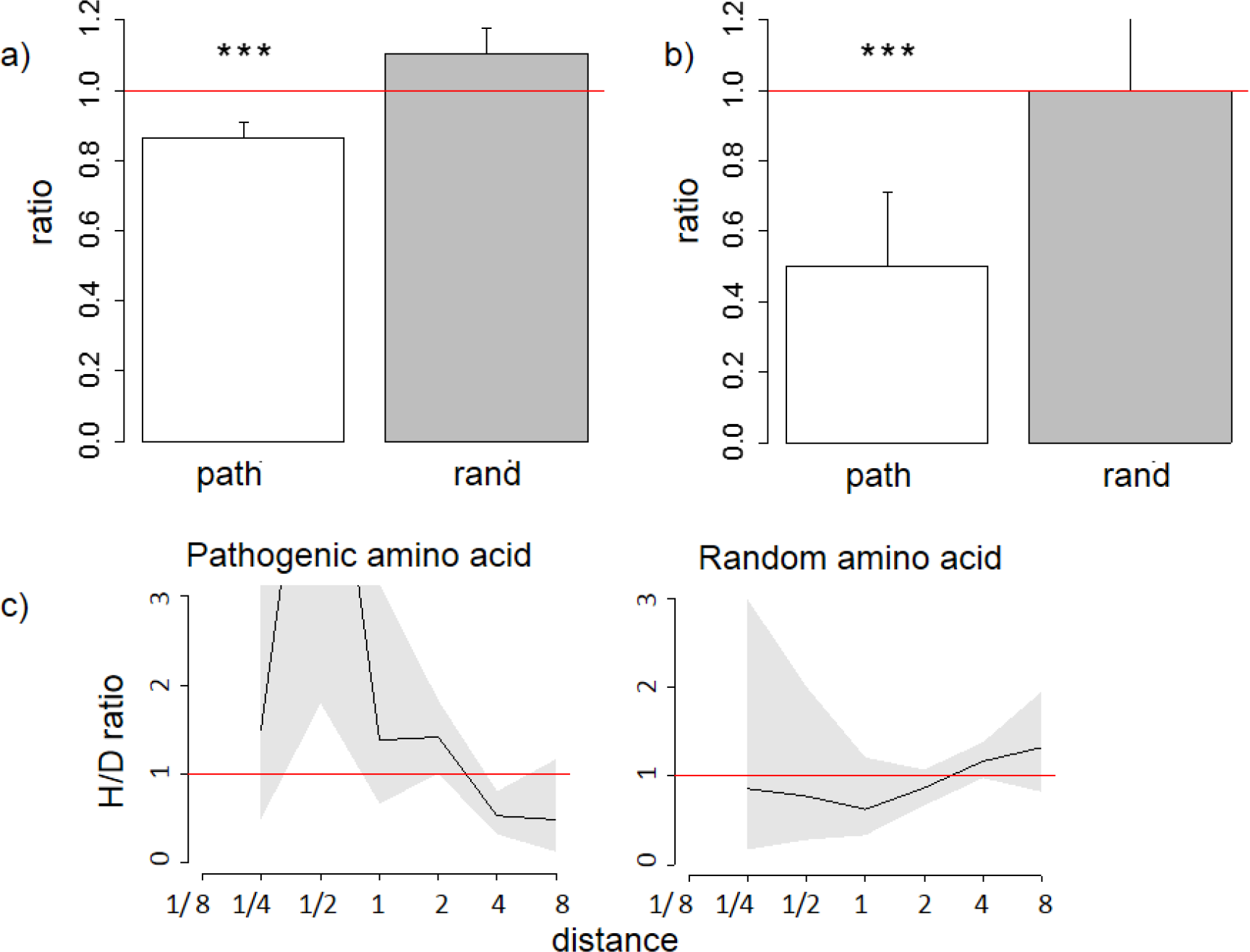
A reduced ratio of the phylogenetic distances between the human branch and substitutions to the considered amino acid vs. to other amino acids at the same site (a,b) and a higher H/D ratio in species closely related to human (c) are observed for all (a,c) and confirmed (b) human pathogenic amino acid, but not for a random allele observed at this site or a simulated allele, in the 4350-species opisthokonts phylogeny. Notations same as in Figures 1 and 2.

### Human pathogenic variants are more biochemically similar than non-human variants to normal human variants

To understand what drives the preferential emergence of the human variant, either normal or pathogenic, in species closely related to humans, we analyzed the identity of these variants. Both normal (Fig. 4a) and pathogenic (Fig. 4b) human amino acids were more similar in their biochemical properties according to the Miyata matrix (Miyata, Miyazawa, & Yasunaga, 1979) to the human reference variant than amino acids observed in non-human species. In turn, amino acids observed in non-human species were more similar to the human reference variant than amino acids never observed at this site in any species.

**Figure 4.**
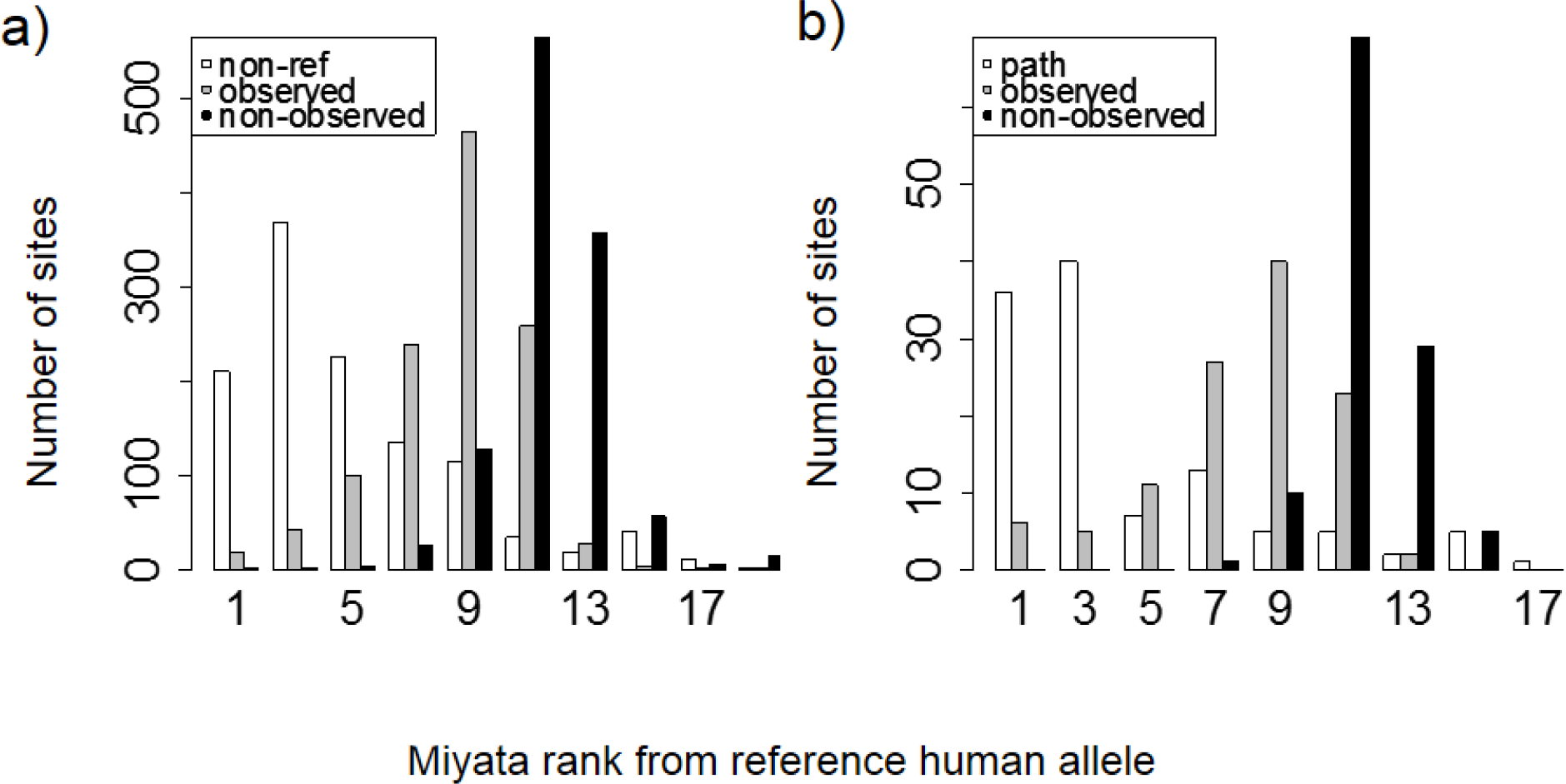
Distributions of ranks of Miyata distances between the reference human allele and the nonreference (a) or pathogenic (b) allele (white), compared with other alleles that were (gray) or were not (black) observed at the same site. For each site with known polymorphisms or pathogenic mutations, we ranked all amino acids by Miyata distance from reference human allele, and then obtained distance rank for pathogenic (or non-reference) human variant, mean rank for amino acids that occurred in a site but did not observed in human and mean rank for rest amino acids.

### Individual mutations

To illustrate the observed phylogenetic clustering, we plotted the distribution over the opisthokont phylogeny of substitutions at the 6 amino acid sites that carry pathogenic mutations with “confirmed” status. Visual inspection of these plots confirms that the substitutions giving rise to pathogenic alleles tend to be clustered in the vicinity of the human, compared to other substitutions of the same ancestral amino acids (SI, figure S9). For the metazoan dataset, we also plot three select individual amino acid sites with known pathogenic mutations which are considered below.

The V→A mutation at ND1 site 113 has been reported to cause bipolar disorder, and decreases the mitochondrial membrane potential and reduces ND1 activity in experiments (Munakata et al., 2004). According to the mtDB database (Ingman & Gyllensten, 2006), the A allele persists in human population at 0.5% frequency. However, we observed that the same allele originated independently in three clades of vertebrates: Old World monkeys (Cercopithecidae), flying lemurs (Cynocephalidae) and turtles (Geoemydidae), while most of the other substitutions of V at this site occurred in invertebrates (figure 5). As a result, the mean phylogenetic distance between human and the parallel V→A substitutions is 2.35 (median 0.75), while it is 4.12 (median 4.6) for substitutions of V to other amino acids.

**Figure 5.**
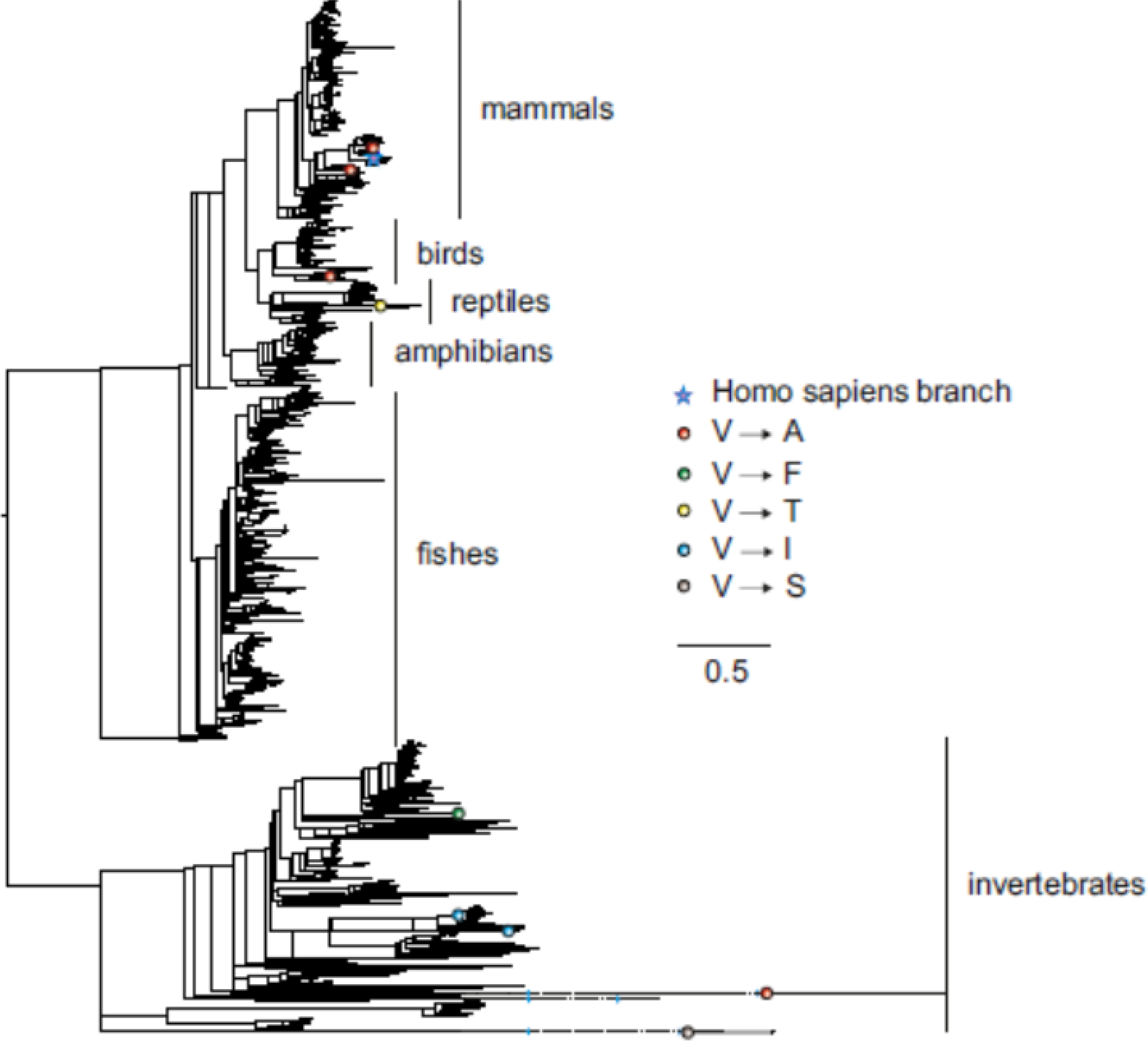
Substitutions in site 113 of ND1. Blue star is *H. sapiens* branch; red dots are substitutions of valine to alanine, which is pathogenic in human and dots of other colors are substitutions to other amino acids. Phylogenetic distances are measured in numbers of amino acid substitutions per site. The branches indicated with the blue waves are shortened approximately by 2 distance units.

The V→A mutation at COX3 site 91 has been reported to cause Leigh disease (Mkaouar-Rebai et al., 2011). In metazoans, A allele at this site has originated independently 7 times, including 6 times from V and once from I. All but one of these substitutions occurred in mammals, while tens of substitutions of V and I giving rise to other amino acids occurred throughout metazoans (Figure 6). As a result, the mean phylogenetic distance from human to V→A substitutions was 0.6 (median 0.7), while the distance to other mutations from V was 1.8 (median 2.1); the corresponding numbers for I were 0.6 (median 0.6) and 2.7 (median 2.2).

**Figure 6.**
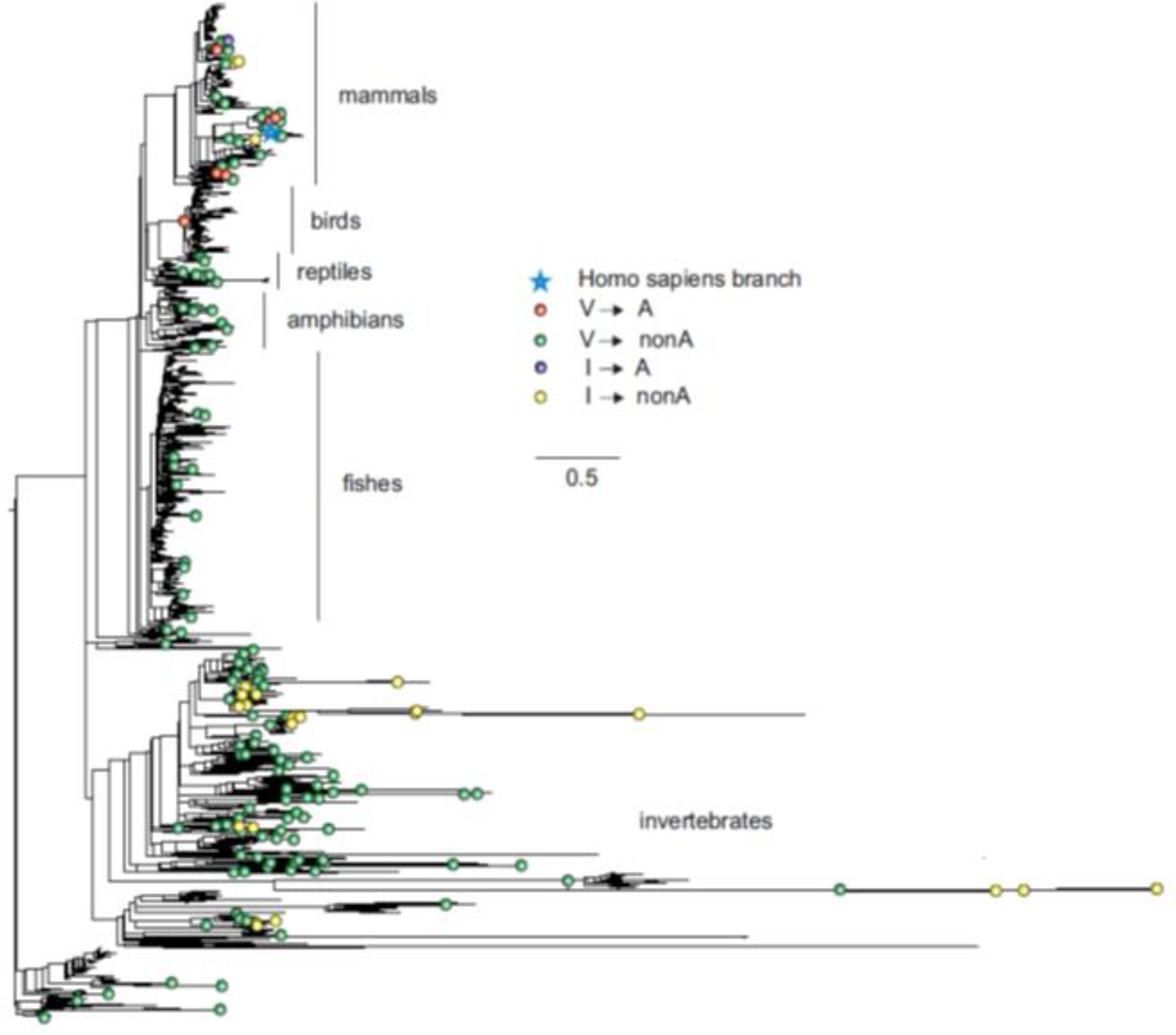
Substitutions in site 91 of COX3. Blue star is *H. sapiens* branch; red dots are substitutions from valine to alanine which is pathogenic in human, black dot is substitution from isoleucine to alanine, and dots of other colors are substitutions to other amino acids. Phylogenetic distances are measured in numbers of amino acid substitutions per site.

To better understand the possible reasons for the unexpected pattern of clustering of the deleterious variant in the phylogenetic vicinity of human, we used molecular dynamics simulations to predict the effect of each mutation on the structure and function of the human protein. As site 91 is positioned within the wall of a pore that is thought to be a channel for oxygen transport (Shinzawa-Itoh et al., 2007), we estimated the pore size that would correspond to each amino acid that occured at this site elswhere on the phylogeny if it arose in the mammalian context. All amino acids led to an increase of the pore size, compared to the normal V allele. Such an increase is expected to permit water molecules to enter the pore, and to impede or prevent oxygen transport. Among the eight observed amino acids, the human pathogenic variant A alters the pore size to the smallest extent, while variants observed in other species significantly increase it, potentially interfering with function (figure 7).

**Figure 7.**
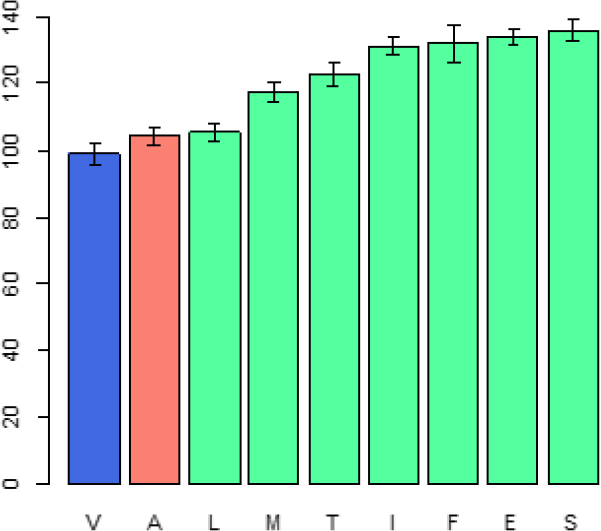
Modelled counts of water molecules in the pore of the human mitochondrial cytochrome c oxidase harboring different mutations in position 91 of COX3. A higher number of molecules probably impedes or prevents oxygen transport. Blue, human reference amino acid (V); red, human pathogenic amino acid (A); green, amino acids observed in non-human species.

Finally, the I→V mutation at ND6 site 33 has been reported to cause type two diabetes (Tawata et al., 2000) and has population frequency of 0.1% according to the mtDB. However, this substitution has occurred repeatedly in parallel in mammals and amphibians, while other substitutions of I were frequent in invertebrates (figure 8). The mean phylogenetic distance from human to parallel substitutions to V was 2.4 (median 1.5), while it was as high as 12.9 (median 14.8) for other amino acids.

**Figure 8.**
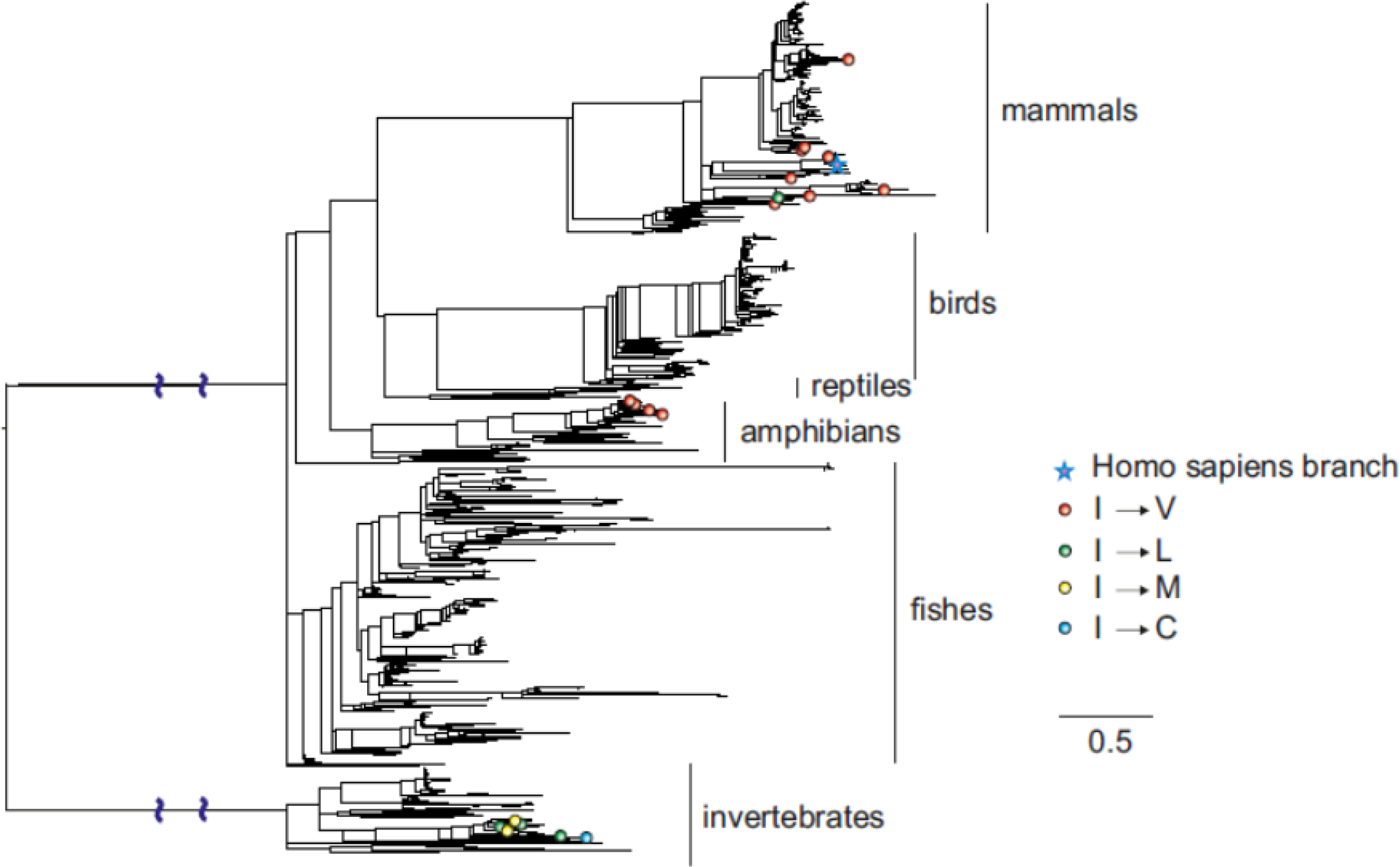
Substitutions in site 33 of ND6. Blue star is *H. sapiens* branch; red dots are substitutions of isoleucine to valine, which is pathogenic in human and dots of other colors are substitutions to other amino acids. Phylogenetic distances are measured in numbers of amino acid substitutions per site. The branches indicated with the blue waves are shortened approximately by 3 distance units.

## Discussion

A variant deleterious in human may be fixed in a non-human species, and sometimes this can be explained by compensatory or permissive mutations elsewhere in the genome (Kondrashov et al. 2002, Kern and Kondrashov 2004, Jordan et al. 2015). Here, we reveal the opposite facet of the same phenomenon: a variant that is fixed or polymorphic in human may be deleterious in a non-human species.

Indeed, we find that substitutions giving rise to the human allele occur in species that are more closely related to *H. sapiens* than species in which substitutions to other amino acid occur. While artefactual evidence for excess of parallel substitutions between closely related species may arise from discordance between gene trees and species trees (Mendes, Hahn, & Hahn, 2016), it is unlikely that it causes the observed signal in our analysis. For reference alleles, we have previously shown that phylogenetic clustering in mitochondrial proteins is not due to tree reconstruction errors (Klink & Bazykin, 2017), and mitochondrial genomes do not recombine, which makes other causes of discordance unlikely. For variants polymorphic in humans, artefactual evidence for parallelism could theoretically also arise from a variant that was polymorphic in the last common ancestor of human and another species such as chimpanzee, was subsequently fixed in this other species, and survived as polymorphism in the human lineage until today. However, the last common ancestor of human mitochondrial lineages probably lived no longer than 148 thousand years ago (Poznik et al., 2013), which is much more recent than the time of human-chimpanzee divergence, also excluding this option.

Therefore, the observed phylogenetic clustering implies a decrease of the fitness conferred by the human amino acid relative to that conferred by other amino acids with phylogenetic distance from human.

Arguably, one would expect the opposite pattern in the variants pathogenic for humans. Indeed, it is likely that the majority of such variants are also deleterious in species related to humans, while changes in the genomic context in more distantly related species may make these variants tolerable.

Instead, we observe the pattern similar to that for the non-pathogenic variants: the amino acid variants pathogenic for humans are also relatively more likely to emerge independently as a homoplasy in species closely related to *H. sapiens* than in more distantly related species.

The similarity between the phylogenetic distribution of the benign and pathogenic variants could be due to some of the benign mutations being incorrectly annotated as pathogenic (Exome Aggregation Consortium et al., 2015). However, none of the six mutations with “confirmed” status in Mitomap are present in the mtDB database among the 2704 sequences from different human populations, suggesting that these mutations are indeed damaging, while they demonstrate a pronounced clustering (figure 3b).

Consideration of biochemical similarities of amino acid variants helps explain why human-pathogenic variants can still be more likely to occur in species closely related to humans. We find that despite their pathogenicity, the human pathogenic variants are on average more biochemically similar to the major human allele than other amino acids that were observed at this site in non-human species. Therefore, in the context of the human genome, the annotated human pathogenic variant probably disrupts the protein structure less than an “alien” non-human variant. Conversely, many alien variants that are not observed in humans are likely to be even more deleterious in human than the annotated pathogenic allele, perhaps lethal, while they confer high fitness in the genomic context of their own species.

Amino acid at position 91 of the COX3 protein provides a case in point. When the mammalian protein structure is used for modelling, we predict that the human pathogenic variant disrupts structure to a smaller extent than each of the seven variants found in reference genomes of other species.

In summary, we have shown that all types of alleles, including pathogenic, that occur in human mitochondrial protein-coding genes more frequently emerge independently in species more closely related to *H. sapiens*. Such a decrease in occurrence of human amino acids with phylogenetic distance from human is probably due to a higher similarity of the sequence and/or environmental context in more closely related species. More generally, it is broadly accepted that the occurrence of a mutation in another species is an important predictor of its pathogenicity in humans (Adzhubei et al., 2010; Kumar, Dudley, Filipski, & Liu, 2011), and it is increasingly appreciated that it is important to account for the degree of relatedness of the considered species to human (Jordan et al., 2015). Our observation that human-pathogenic alleles are underrepresented in more distantly related, rather than in more closely related, species shows that the direction of the association between relatedness and prediction of pathogenicity can be counterintuitive.

## Acknowledgements

We thank Shamil Sunyaev, Alexey Kondrashov, Dmitry Pervouchine and Vladimir Seplyarskiy for valuable comments.

---

**Figure S1.**
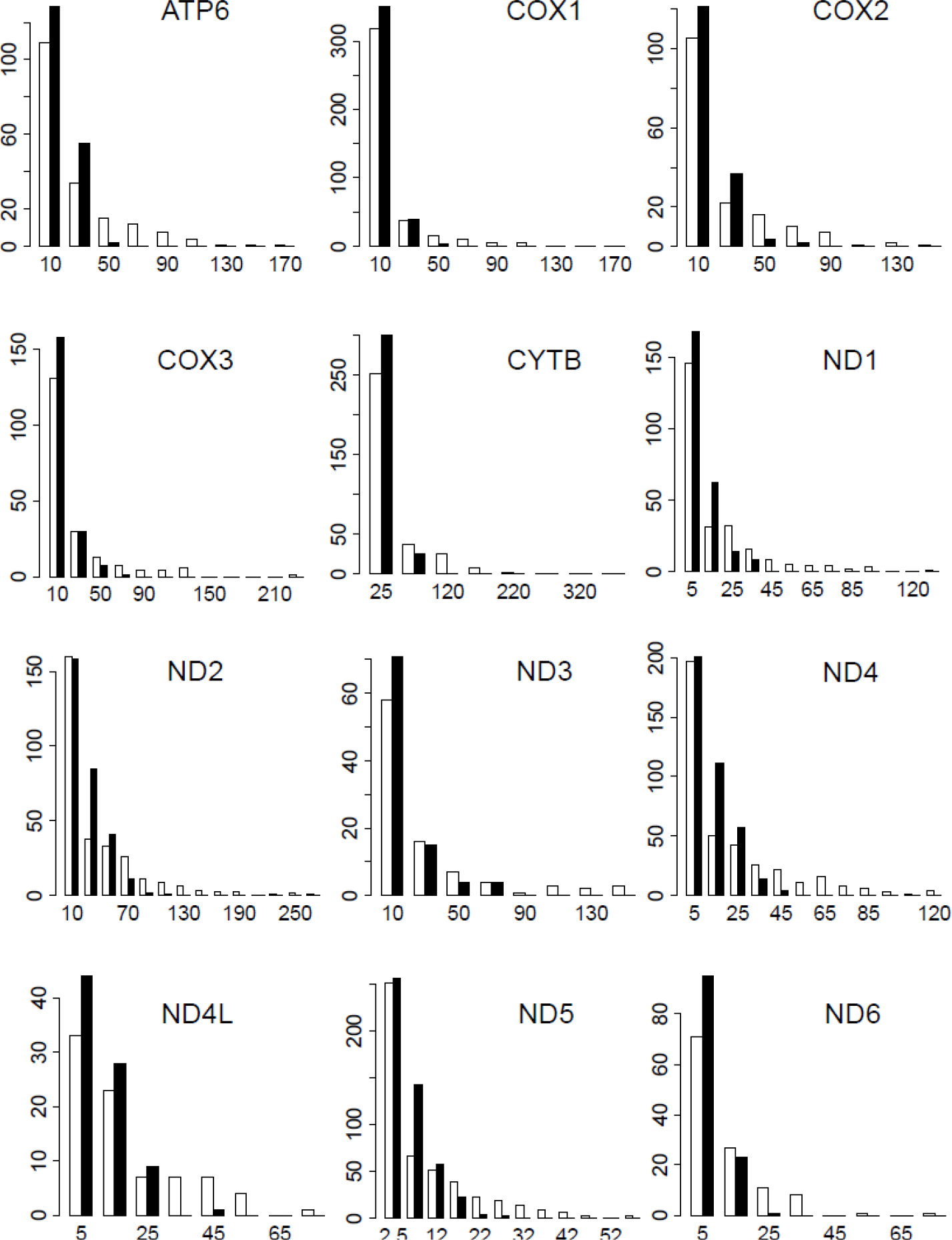
Numbers of substitutions that give rise to the human amino acid variant (white) or the mean number of substitutions to a particular amino acid (black) at a site for sites with at least one substitution in Metazoan dataset.

**Figure S2.**
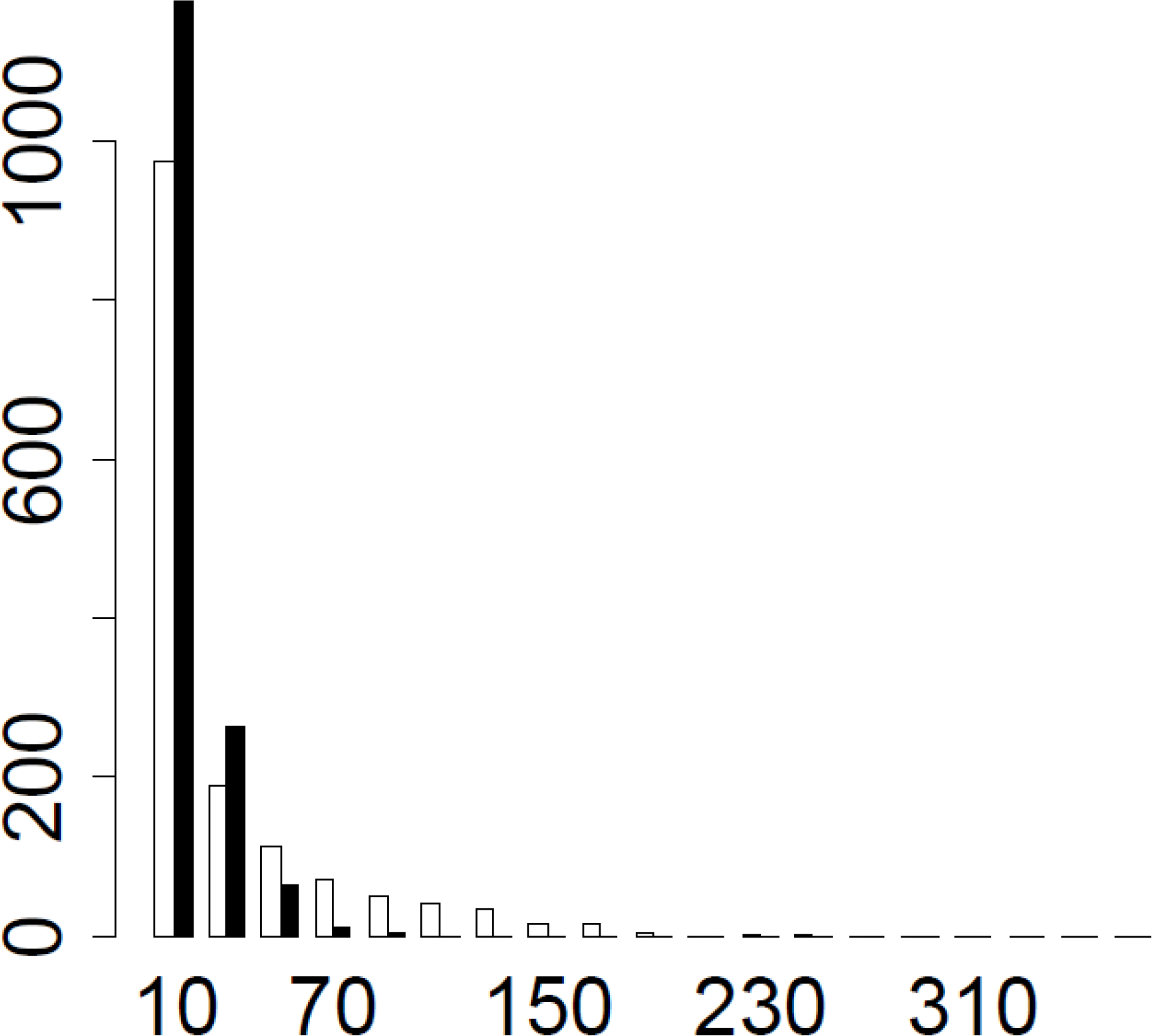
Numbers of substitutions that give rise to the human amino acid variant (white) or the mean number of substitutions to a particular amino acid (black) at a site for sites with at least one substitution in opisthokonts dataset.

**Figure S3.**
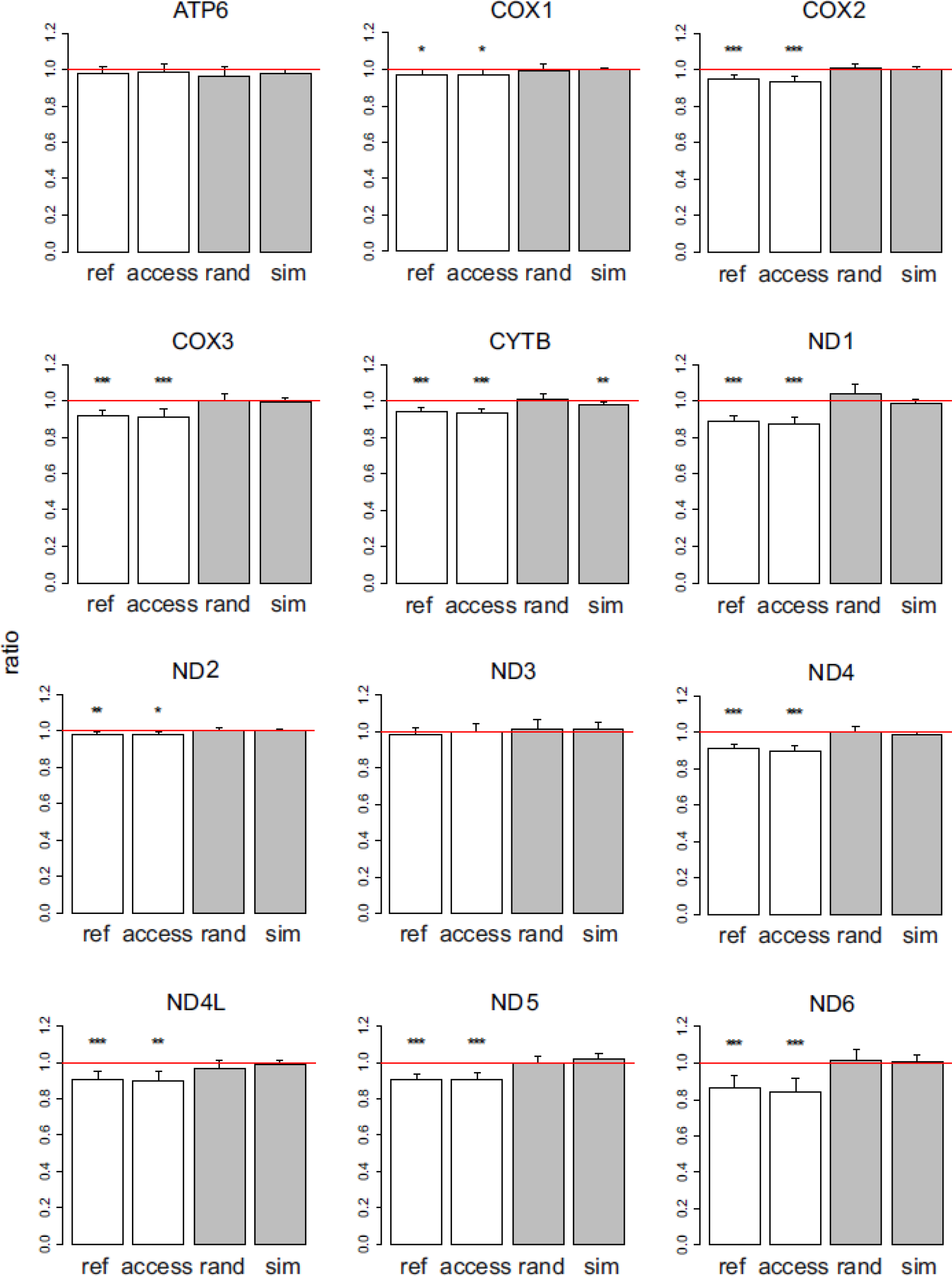
Ratios of the phylogenetic distances between the human branch and substitutions to the human reference allele vs. to other amino acids, for metazoan dataset. Ratios <1 imply that the considered allele arises independently closer at the phylogeny to humans than other alleles. The bar height and the error bars represent respectively the median and the 95% confidence intervals obtained from 1,000 bootstrap replicates, and asterisks show the significance of difference from the one-to-one ratio (*, P<0.05; **, P<0.01;***, P<0.001). ref, human reference allele; access, human reference alleles from accessible amino acid pairs (see text); rand, a random non-human amino acid among those that were present in the site; sim, human allele in simulated data.

**Figure S4.**
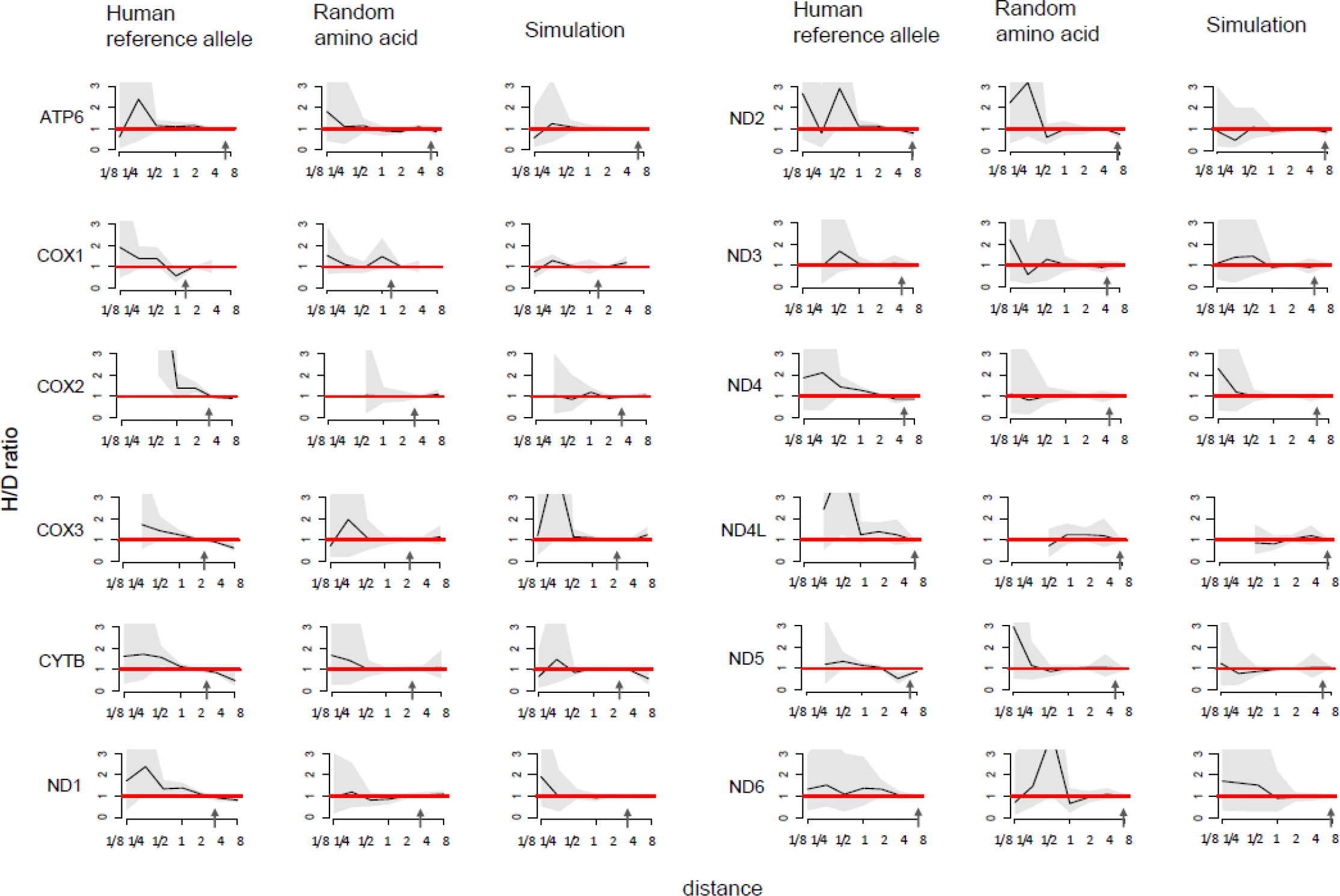
Higher fraction of homoplasic substitutions to the human reference amino acid, compared with random amino acids that had independently originated at this site (H/D ratio), in species closely related to human, for metazoan dataset. Horizontal axis, distance between branches carrying the substitutions and the human branch, measured in numbers of amino acid substitutions per site, split into bins by log_2_(distance). Vertical axis, H/D ratios for substitutions at this distance. Black line, mean; grey confidence band, 95% confidence interval obtained from 1000 bootstrapping replicates. The red line shows the expected H/D ratio of 1. Arrows represent the distance between human and *Drosophila*.

**Figure S5.**
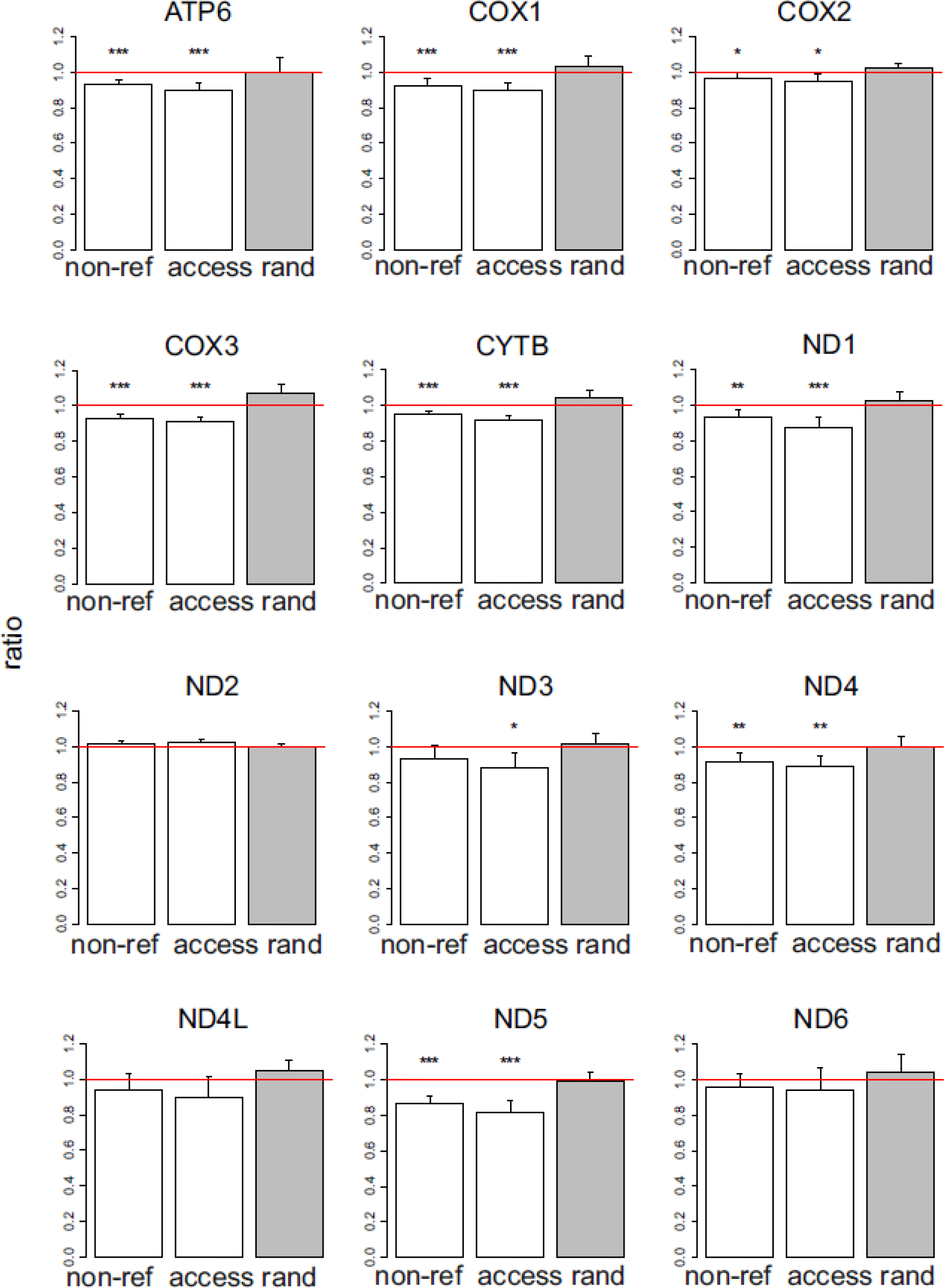
Ratios of the phylogenetic distances between the human branch and substitutions to the human non-reference allele vs. to other amino acids, for metazoan dataset. Notations same as in Figure S1.

**Figure S6.**
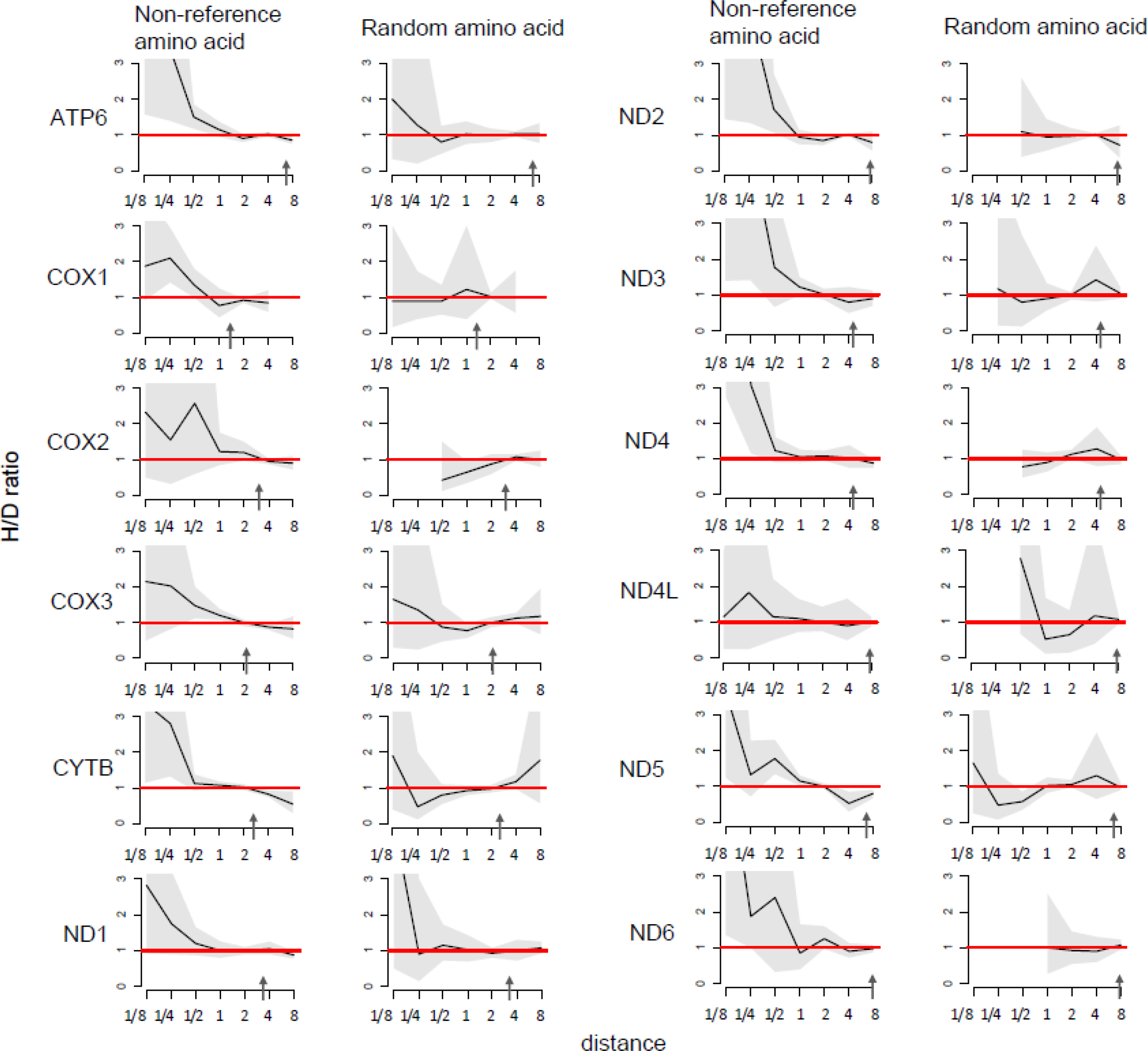
Higher fraction of homoplasic substitutions to the human non-reference amino acid, compared with random amino acids that had independently originated at this site (H/D ratio), in species closely related to human, for metazoan dataset. Notations same as in Figure S2.

**Figure S7.**
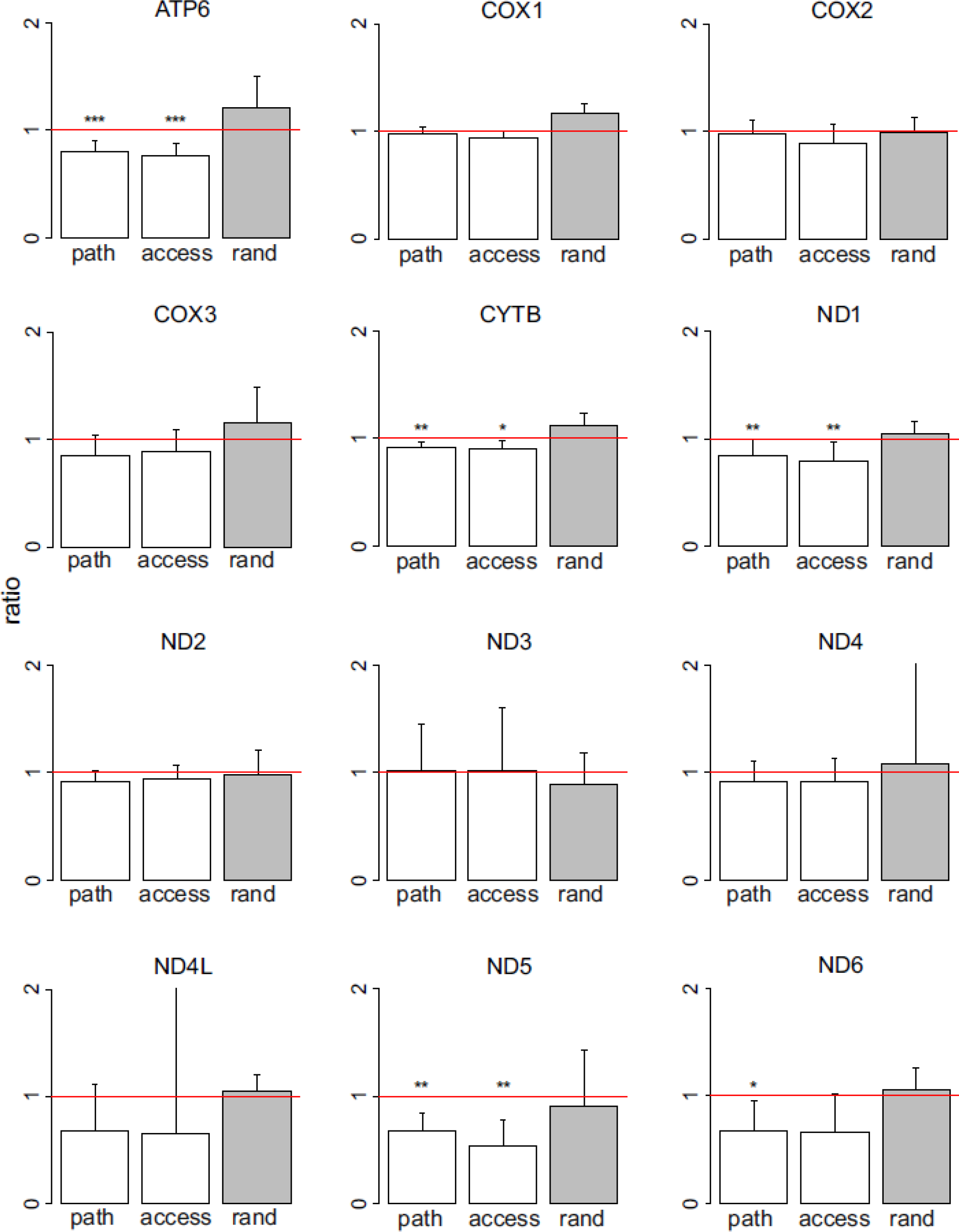
Ratios of the phylogenetic distances between the human branch and substitutions to the human pathogenic allele vs. to other amino acids, for metazoan dataset. Notations same as in Figure S1.

**Figure S8.**
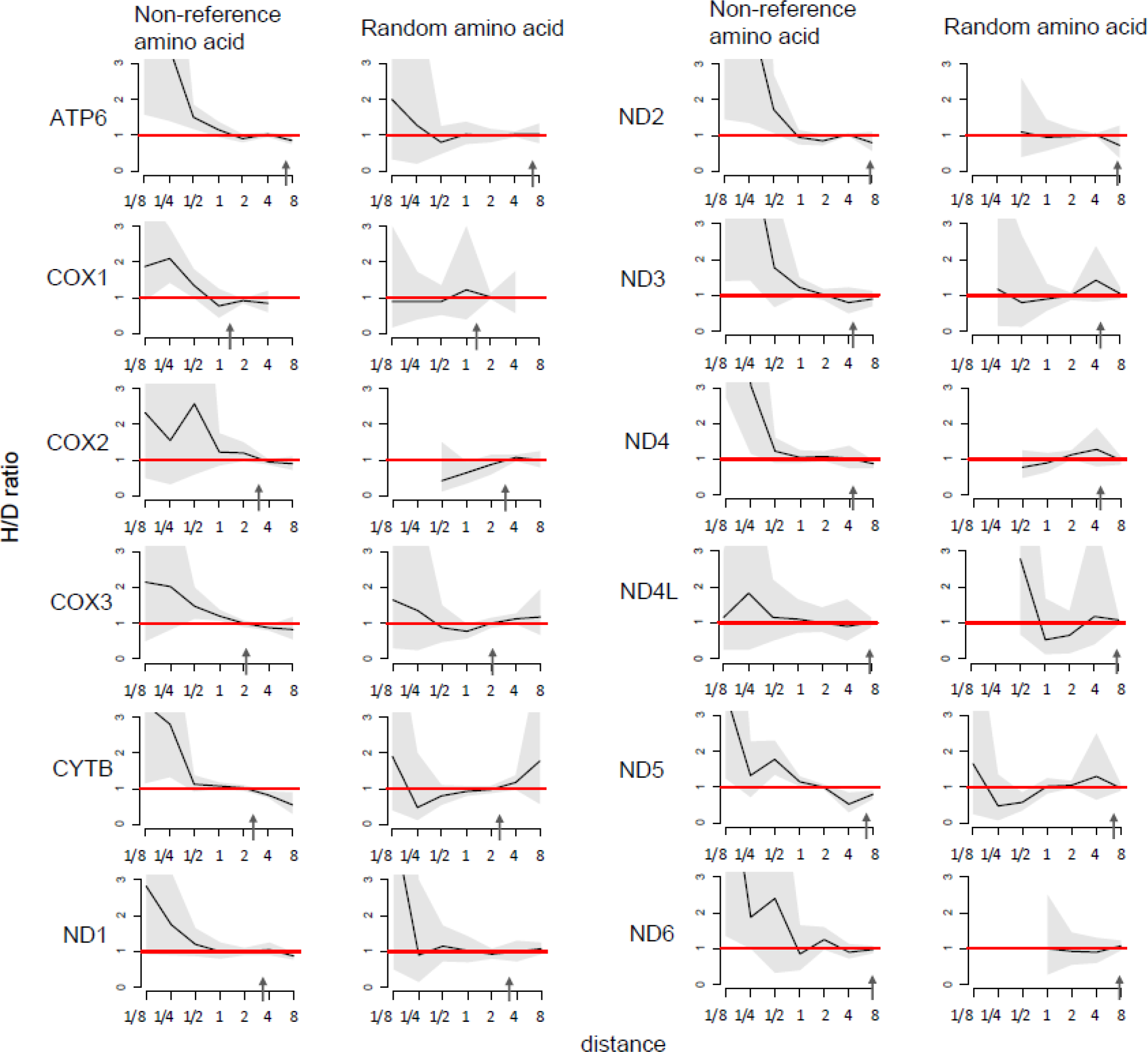
Higher fraction of homoplasic substitutions to the human pathogenic amino acid, compared with random amino acids that had independently originated at this site (H/D ratio), in species closely related to human, for metazoan dataset. Notations same as in Figure S2.

**Figure S9.**
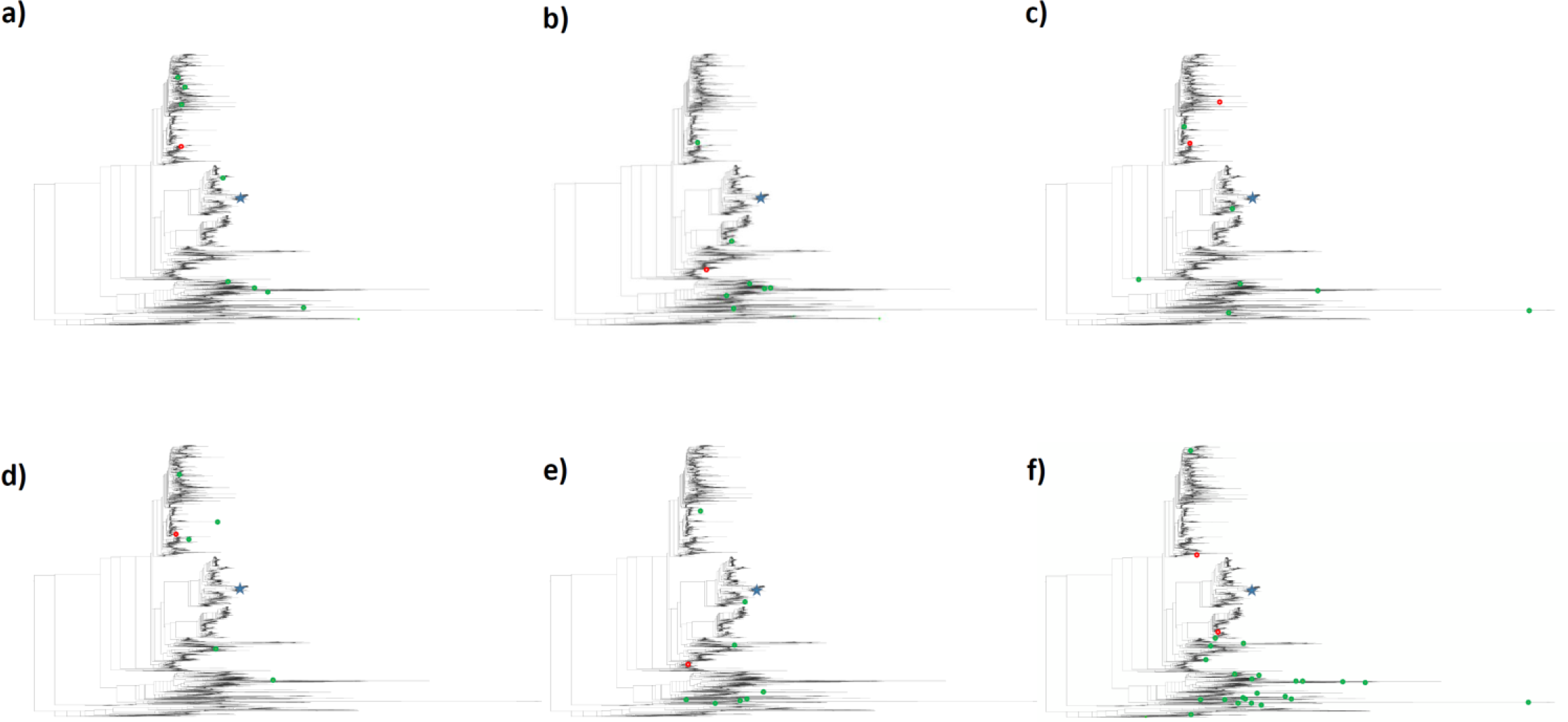
Substitutions giving rise to the human pathogenic allele (red circles) and to other amino acids (green circles) on the 4350-species phylogeny. a, site 156 of ATP6; b, site 217 of ATP6; c, site 220 of ATP6; d, site 278 of CYTB; e, site 35 of CYTB; f, site 40 of CYTB. Blue star, *H. sapiens*.

**supplementary Table 1.** Homoplasies giving rise to the human reference allele.

| **Gene** | **Species** | **Sites** | **Analyzed sites** | **Substitutions to the human amino acid per site** | **Substitutions to the human amino acid per site in simulation** |
| --- | --- | --- | --- | --- | --- |
| ATP6 | Metazoans | 186 | 131 | 25.3 | 16.7 |
| COX1 | Metazoans | 404 | 146 | 12.1 | 13.9 |
| COX2 | Metazoans | 165 | 110 | 22.5 | 22.3 |
| COX3 | Metazoans | 198 | 128 | 22.4 | 16.2 |
| CYTB | Metazoans | 327 | 183 | 32.2 | 29.7 |
| ND1 | Metazoans | 253 | 160 | 15.5 | 12.1 |
| ND2 | Metazoans | 299 | 220 | 36.3 | 34.0 |
| ND3 | Metazoans | 94 | 67 | 26.6 | 20.0 |
| ND4 | Metazoans | 392 | 263 | 19.9 | 17.3 |
| ND4L | Metazoans | 82 | 70 | 18.4 | 19.6 |
| ND5 | Metazoans | 516 | 302 | 8.7 | 7.3 |
| ND6 | Metazoans | 119 | 91 | 11.4 | 10.1 |
| ATP6+COX1+COX2+COX3+CYTB | Opisthokonts | 1524 | 964 | 30.2 | 24.4 |

**supplementary Table 2.** Homoplasies giving rise to the human non-reference allele.

| **Gene** | **Species** | **Polymorphic sites** | **Analyzed sites** | **Substitutions to the human non-reference allele per non-reference allele** | **Mean no. of non-reference alleles per polymorphic site** |
| --- | --- | --- | --- | --- | --- |
| ATP6 | Metazoans | 142 | 12 | 21.1 | 1.9 |
| COX1 | Metazoans | 131 | 10 | 21.0 | 1.4 |
| COX2 | Metazoans | 81 | 72 | 28.3 | 1.4 |
| COX3 | Metazoans | 108 | 93 | 24.8 | 1.6 |
| CYTB | Metazoans | 199 | 18 | 36.0 | 1.7 |
| ND1 | Metazoans | 101 | 88 | 16.8 | 1.5 |
| ND2 | Metazoans | 143 | 64 | 49.1 | 1.6 |
| ND3 | Metazoans | 34 | 32 | 30.3 | 1.6 |
| ND4 | Metazoans | 136 | 10 | 20.9 | 1.2 |
| ND4L | Metazoans | 29 | 26 | 17.3 | 1.5 |
| ND5 | Metazoans | 223 | 17 | 9.9 | 1.6 |
| ND6 | Metazoans | 55 | 45 | 11.1 | 1.5 |
| ATP6+COX1+COX2+COX3+CYTB | Opisthokonts | 775 | 516 | 32.2 | 1.6 |

**supplementary Table 3.** Homoplasies giving rise to the human pathogenic variants.

| Gene | Species | Sites with pathogenic alleles | Analyzed sites | Substitutions to the human pathogenic amino acid per pathogenic amino acid | Mean no. of pathogenic amino acids per site with pathogenic alleles |
| --- | --- | --- | --- | --- | --- |
| ATP6 | Metazoans | 15 | 11 | 9.7 | 1.1 |
| COX1 | Metazoans | 20 | 15 | 41.0 | 1.0 |
| COX2 | Metazoans | 10 | 9 | 44.8 | 1.0 |
| COX3 | Metazoans | 9 | 7 | 14.8 | 1.0 |
| CYTB | Metazoans | 20 | 16 | 15.5 | 1.1 |
| ND1 | Metazoans | 25 | 17 | 6.3 | 1.1 |
| ND2 | Metazoans | 10 | 8 | 29.5 | 1.0 |
| ND3 | Metazoans | 6 | 5 | 3.5 | 1.0 |
| ND4 | Metazoans | 7 | 4 | 15.1 | 1.2 |
| ND4L | Metazoans | 3 | 3 | 6.0 | 1.0 |
| ND5 | Metazoans | 24 | 8 | 5.1 | 1.0 |
| ND6 | Metazoans | 13 | 10 | 8.0 | 1.1 |
| ATP6+COX1+COX2+ COX3+CYTB | Opisthokonts | 89 | 72 | 25.0 | 1.0 |

